# Multi-omics phenotyping characterizes molecular divergence underlying different clinical scenarios of inflammatory bowel disease

**DOI:** 10.1101/2024.05.13.593901

**Authors:** Nguyen Tran Nam Tien, Eun Jeong Choi, Nguyen Quang Thu, Seung Jung Yu, Duc Ninh Nguyen, Dong Hyun Kim, Nguyen Phuoc Long, Hong Sub Lee

## Abstract

Clinically heterogeneous spectrum and molecular phenotypes of inflammatory bowel disease (IBD) remain to be comprehensively elucidated. This study set out to explore the serum molecular profiles (I) of IBD subtypes; in association with (II) elevated fecal calprotectin and (III) disease activity states; (IV) upon treatment escalation; and (V) in patients who needed treatment escalation. The serum proteome, metabolome, and lipidome of 75 treated IBD patients were profiled. Single- and multi-omic data analysis was performed to determine differential analytes and integrative biosignatures. (I) Chronic inflammation, and phosphatidylcholine and bile acid homeostasis disturbances underlined the differences between Crohn’s disease (CD) and ulcerative colitis. (II) Elevated calprotectin was associated with higher levels of inflammatory proteins and sphingomyelins (SM) and lower levels of bile acids, amino acids, and triacylglycerols (TG). Relative to patient remission, active disease state (III) was characterized by decreased SMs and increased inflammatory proteins and TGs. (IV) Treatment escalation was associated with augmented levels of inflammatory response-related proteins and reduced levels of amino acids. Most TG species increased in the post-treatment escalation. Moreover, needed-treatment-escalation patients had significantly lower levels of TGs (V). They also showed increased SMs and decreased signaling receptor binding proteins. Multi-omics analysis revealed biosignatures that captured the differences between groups of each scenario. Eight analytes, including NFASC, ANGPTL4, and chenodeoxycholate, were found in at least three biosignatures. Collectively, disturbances in immune response, bile acid homeostasis, amino acids, and lipids alteration potentially underlie the clinically heterogeneous spectrum of IBD.

## INTRODUCTION

Inflammatory bowel disease (IBD) poses a significant health burden worldwide, impacting millions of individuals ^1,2^. The disease emerges from a multifaceted interplay involving genetic predisposition, the gut microbiota, diverse environmental exposures, and immunological disturbances ^3,4^. Upon diagnosis, IBD is typically classified as Crohn’s disease (CD) and ulcerative colitis (UC) ^5^. Treatment plans are tailored based on disease activity, aiming to mitigate inflammation, alleviate symptoms, and enhance the patient’s quality of life ^6^. Multiple evaluation methods are needed to achieve this goal.

Biomarkers including fecal calprotectin serve as a surrogate endpoint of gastrointestinal tract inflammation and clinicians implement symptomology to manage the patients ^6^. Failure of standard therapy requires treatment escalation where biologics target key immunoregulatory-related processes to reduce inflammation, improve symptoms, and promote healing for IBD patients ^7,8^. However, disease heterogeneity, inadequate drug concentration, and immunogenicity development can compromise biologics’s efficacy ^4^. Therefore, a deeper understanding of molecular profiles associated with the disease pathogenesis, inflammation level, disease activity state, and molecular profiles upon treatment escalation is essential ^5^.

Deep molecular phenotyping represents a paradigm shift in disease characterization and refers to a comprehensive and highly detailed approach beyond solely relying on clinical, biochemical, and imaging findings ^9,10^. Multi-omics stands out as a promising approach for comprehensively characterizing complex diseases like IBD. Indeed, the approach has gradually become a cornerstone of IBD research ^11–19^. Several molecular perturbations driving disease pathophysiology, including barrier dysfunction, chronic immunological response, microbial dysbiosis, and consequences alteration in tricarboxylic acid cycle metabolites, amino acid, lipid metabolism, and oxidative pathways, have been elucidated thanks to the omics approach. However, gaps persist in current IBD omics-based research, e.g., limited independent validation and a small number of multi-omic studies ^18^. Additionally, most (multi)-omics studies focused on molecular profiling for IBD subtypes, with a limited number of studies for other clinical scenarios, including disease activity, inflammation level, and treatment response.

This study aimed to explore the differences in serum molecular profiles (proteome, metabolome, and lipidome) underpinning different clinical scenarios of the under-treatment IBD population. Specifically, we characterized the single- and multi-ome differences among patients diagnosed with either CD or UC, those with elevated or normal fecal calprotectin levels, and individuals who presented an active or remission disease state. Moreover, we investigated the molecular perturbations in patients who underwent treatment escalation with samples collected before and after the treatment initiation, as well as those who needed treatment escalation and those who did not. The study provided comprehensive insights into the functional multi-ome profiles of IBD patients across five distinct clinical scenarios and laid a foundation for subsequent studies on improving disease characterization and management.

## METHODS

### Ethical statement and study population

The study protocol was reviewed and approved by the Institutional Review Board of Busan Paik Hospital (No. BPIRB 2024-02-046). The study was conducted following the Helsinki Declaration. Before enrollment, all subjects provided written informed consent. Previously diagnosed IBD patients who underwent extensive treatment were recruited in the current study. Participants were excluded if they had severe comorbidities or extraintestinal manifestations. Endoscopy was performed to classify patients into UC and CD subtypes at enrolment (**Table 1**). Fecal calprotectin was used to categorize participants into elevated or normal groups using a 150 µg/g cut-off ^20,21^. Median imputation was used in 2 cases with missing calprotectin levels. The C-reactive proteins were significantly higher in the CD than in the UC group and in the elevated fecal calprotectin group compared to the normal counterpart (**Table 1**, **Supplementary Table S1**). The MAYO score cut-off of 1 and CD activity index (CDAI) score cut-off of 150 concerning CD and UC were used to reflect the disease status as either active state (i.e., mild, moderate, or severe conditions) or remission. Remission-state patients tended to have a more prolonged illness duration (**Supplementary Table S2**). Patients with samples collected before and after the treatment escalation were assigned to the pseudo-pre- and post-treatment escalation design as an unpaired comparison (**Supplementary Table S3**). The former group (i.e., pre-treatment escalation) also served as patients who needed treatment escalation, compared to those who did not need it (**Supplementary Table S4**). These constituted five clinically meaningful scenarios, which are schematized in **Fig. 1**.

**Fig. 1.**
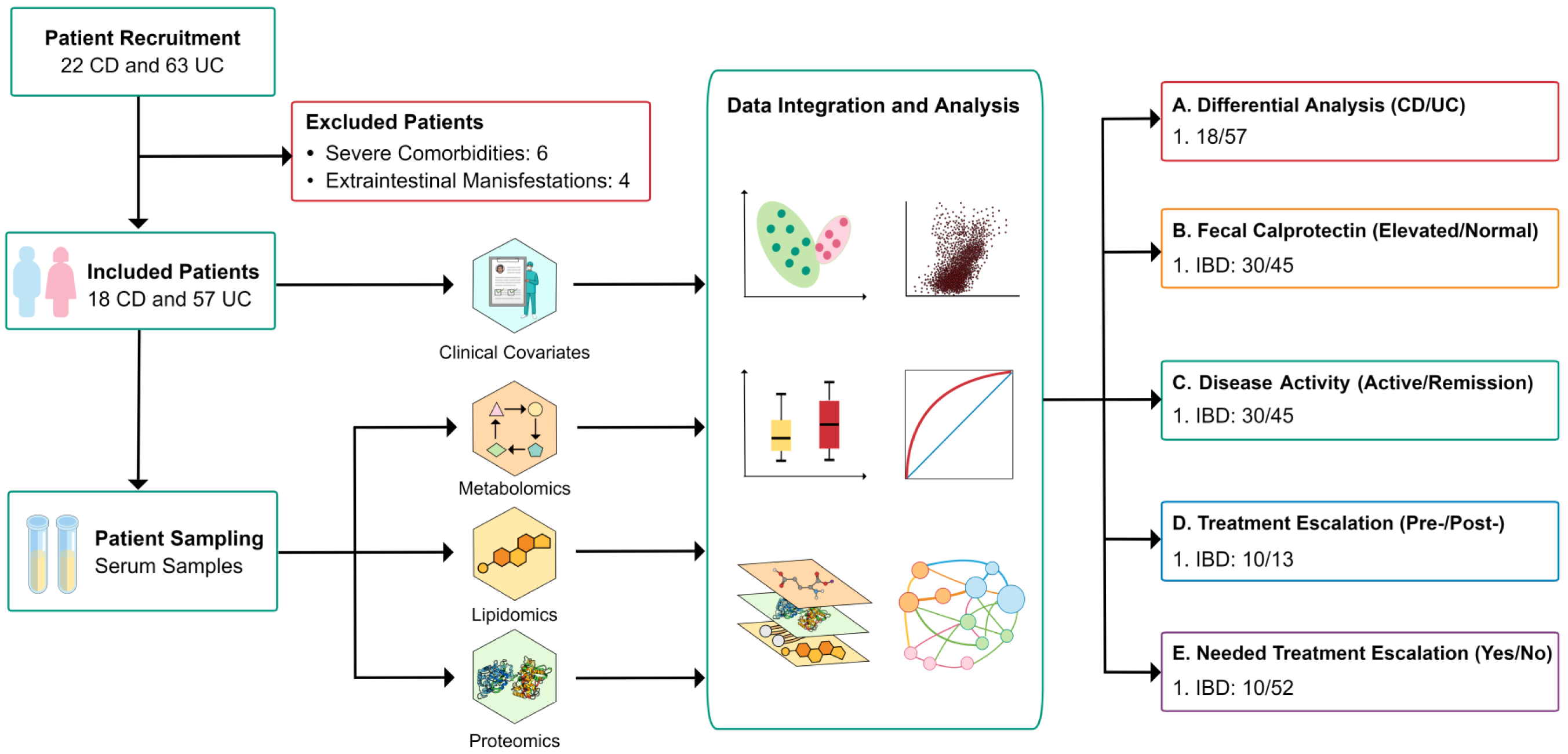
Overall study workflow spanning from patient recruitment, sampling, multi-ome profiling, and data integration, analysis, and interpretation of five clinically meaningful scenarios of treated inflammatory bowel disease patients.

**Table 1.**
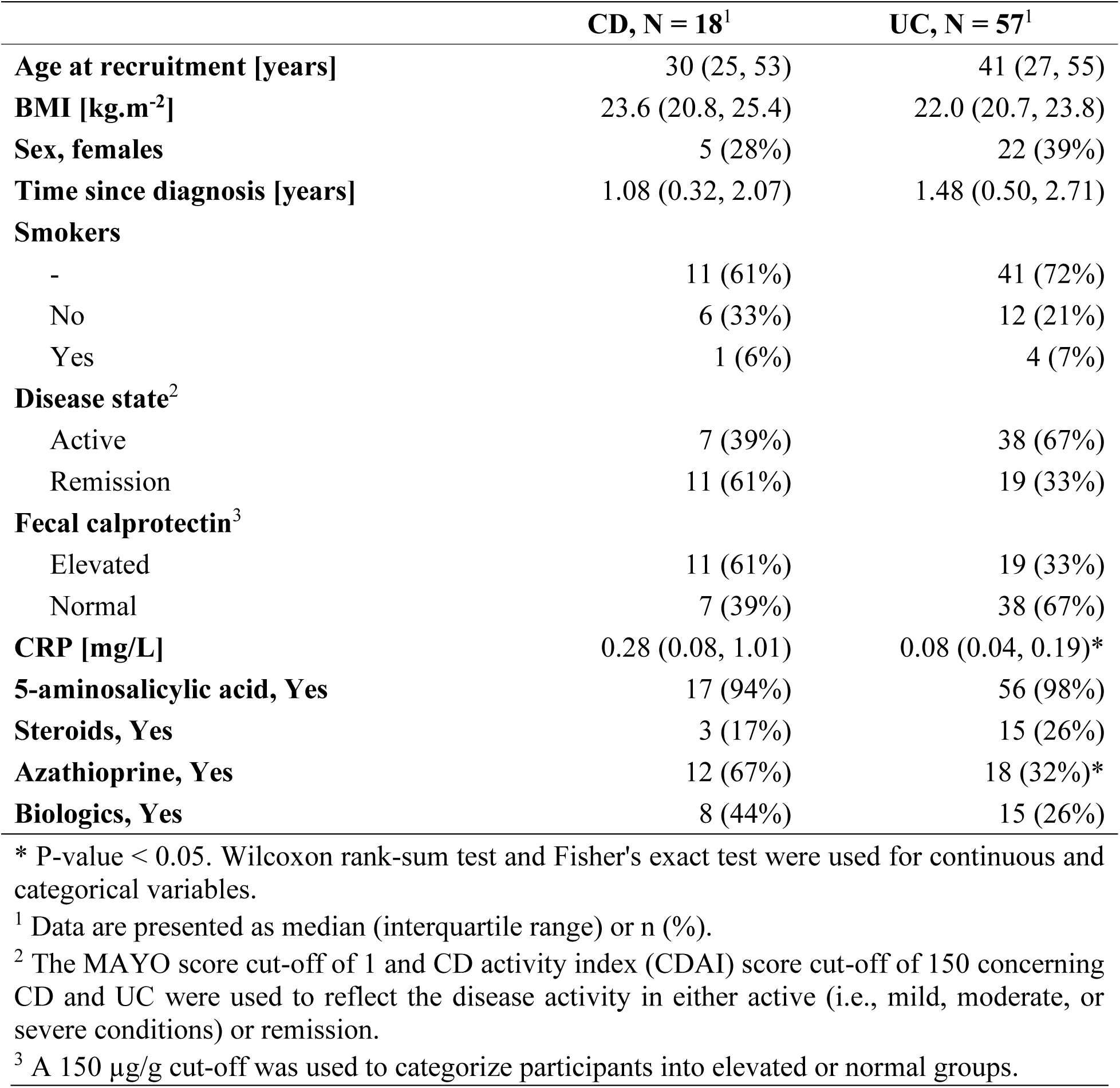
Clinical characteristics by disease subtypes.

### Metabolomics and lipidomics sample preparation and data acquisition

The metabolomics and lipidomics experiments followed our previously established procedure ^22–26^. Detailed plasma sample preparation, sample treatment, and instrumental analysis of metabolomics and lipidomics were described in the **supplementary methods**.

### Proteomics sample preparation and data acquisition

For proteomics analysis, Olink® panels of Explore 384 Inflammation and Target 96 Cardiovascular III (Olink, Uppsala, Sweden) were used. Sample treatment for protein extraction followed Olink’s established procedure. This method utilizes a highly specific antibody probe labeled with two oligonucleotides to bind the target protein. The real-time PCR amplification of the oligonucleotide sequence then allows for quantitative measurement of the resulting DNA sequence. The experiment was conducted by DNALink (Seoul, Republic of Korea).

### Data preprocessing, data treatment, and normalization

MS-DIAL (version 4.9.0) was used for MS data processing ^27^. In-house libraries of authentic standards, MS-DIAL in-built libraries, and public libraries were used to annotate metabolites and lipids ^23,24^. Only metabolites and lipids with reliable annotations were subjected to subsequent analyses. Next, the aligned data was imported to the *MetaboAnalystR* package version 4.0.0 for further data treatment ^28^. The normalized protein expression values were used for proteomics data, and the values recorded below the limit of detection were considered missing. Data points from the Olink® Target 96 Cardiovascular III were used for downstream analyses when proteins were quantified in 2 panels.

Analytes (metabolites, lipids, and proteins) with a missing rate ≥ 50% were excluded, whereas the retained missing values were inputted via the k-nearest neighbors algorithm. For metabolomics and lipidomics, analytes whose relative standard deviation was > 25% in the pooled QCs samples were also filtered out.

### Single-omics differential analysis

For differential analysis, the *limma* method was used. The analysis was performed in the *MetaboAnalystR* package version 4.0.0 for each omics data separately ^28^. Pre-treated and normalized data without scaling was used to obtain differential analytes (DAs) of metabolites, lipids, and proteins between groups. A P-value of 0.05 was considered the putative significant cut-off. The heatmap of DAs of each omics was visualized via the *ComplexHeatmap* version 2.14.0 in R 4.2.3 ^29^.

### Integrative multi-omics analysis and biosignature identification

Regarding integrative multi-omics analysis, we employed the data integration analysis for biomarker discovery using the latent components (DIABLO) method, which was performed using the *mixOmics* package version 6.23.4 ^30,31^. Normalized data of each omics data was used as input data. The scaling was conducted internally by DIABLO. For model tuning, we first explored the weight and chose the value of 0.5 to satisfy a trade-off between correlation and discriminability. Next, the number of components and the distance method were selected via a 10-fold cross-validation (CV). Afterward, the number of features within each data layer was determined. The performance of selected biosignatures to discriminate groups of interest was obtained separately. The area under the receiver operating characteristic curve (AUC) was calculated by averaging sequentially over 10-fold cross-validation, 10-repeated, and three data blocks ^30,31^.

## RESULTS

To investigate the molecular perturbation underlying different clinical scenarios of IBD, we stratified the patient cohort using their diagnostic results, fecal calprotectin level, disease activity score, and treatment escalation information (**Fig. 1**).

### Treated UC and CD patients presented divergence in molecular perturbations linked to chronic inflammation, phosphatidylcholines, and bile acid homeostasis

UC and CD are traditionally the two main subtypes of IBD, which are macro-phenotyped based on endoscopy results and clinical manifestations (**Fig. 2a**). Herein, we determined molecules had differential abundance between UC *vs.* CD. Employing an age- and sex-adjusted linear model, we found 44, 11, and 29 differential proteins, metabolites, and lipids, respectively. These differential analytes (DAs) showed relatively straightforward patterns of differences across three data modalities, especially lipids, and proteins (**Fig. 2b-d**). Most differential proteins (e.g., IFNG, IL1R1, IL12RB1, CCL20, AGER) were linked to inflammation and immune signaling pathways. We also observed a consistent up-regulation of cell adhesion molecules, including ITGB2, ITGB6, ITGA11, LAMA4, and ADA, among others (**Fig. 2b**). Regarding metabolomics modality, changes in bile acid biosynthesis were observed where levels of three conjugated bile acids (glycocholate, taurocholate, and taurodeoxycholate) increased 2-3.5 times in the UC compared to the CD group. Changes of two amino acids (dihydroorotate and 5-hydroxylysine) were also observed (**Fig. 2c**). Lipidomics analyses showed that phosphatidylcholines (PC) and its derivatives of lysoPC (LPC) and ether-linked PC [PC(O-)] and LPC(O-) were the major altered species. Abundances of LPC, lysophosphatidylethanolamines (LPE), ceramides (Cer), and triacylglycerols (TG) were decreased, whereas levels ether-linked phosphatidylethanolamine [PE(O-)] were augmented in the UC group. PCs showed up- and down-regulation patterns (**Fig. 2d**).

**Fig. 2.**
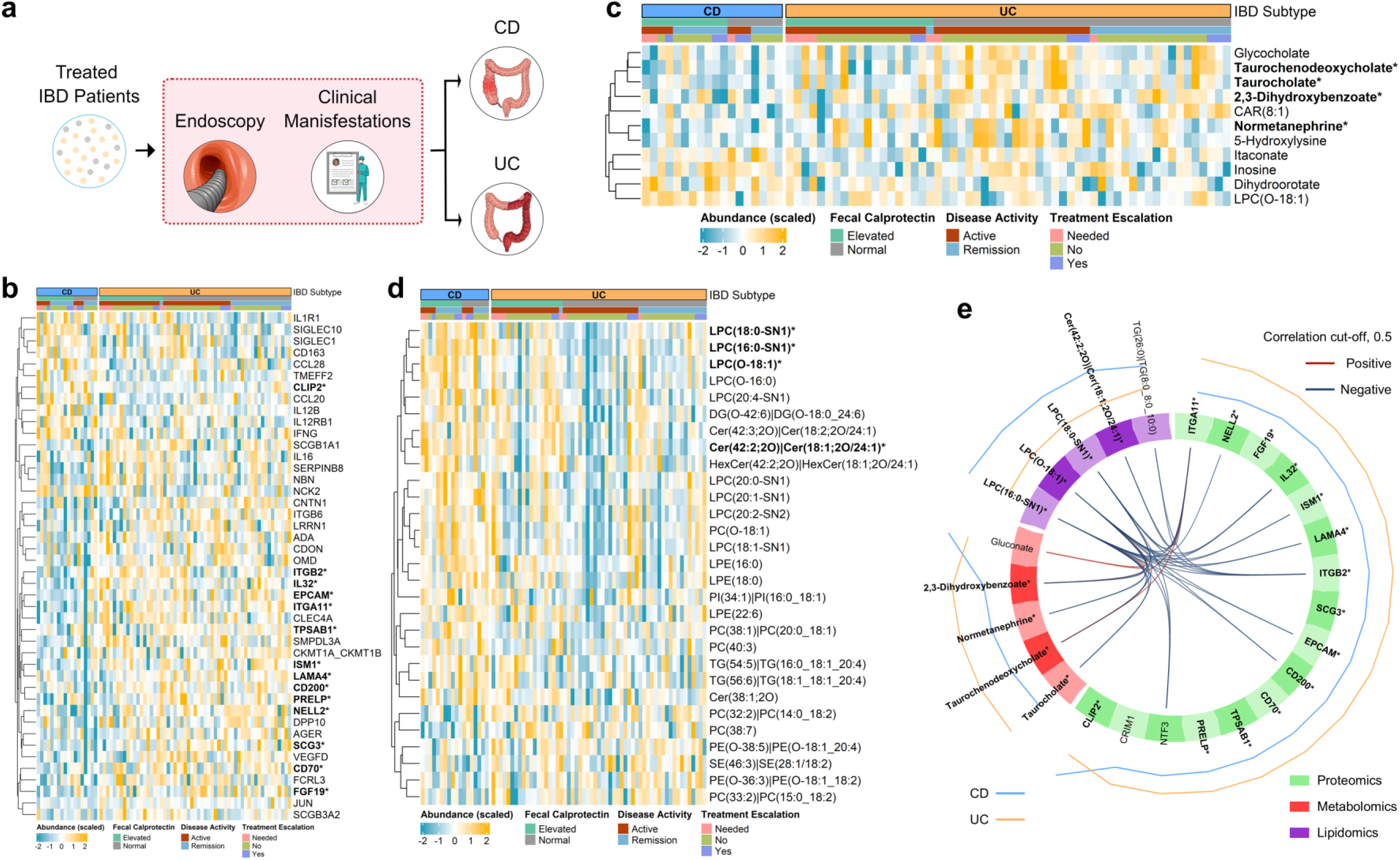
Single and integrative omics analysis presenting molecular perturbances between treated UC and CD patients. **(a)** Sample collection and grouping. Heatmap of differential **(b)** proteins, **(c)** metabolites, and **(d)** lipids. **(e)** DIABLO-selected biosignatures. Abbreviations: IBD: inflammatory bowel disease, UC: ulcerative colitis, CD: Crohn’s disease. * and bold: analytes detected in both single- and multi-omics data analysis.

Utilizing an integrative omics approach, we evaluated the correlation of the data layers and determined the highly correlated and discriminant features across the three data blocks. The proteomics modality correlated well with two others (**Fig. 2e**, **Supplementary Fig. S1a**). Among 16 DIABLO-identified proteins, most of them showed significantly higher abundance in UC, except an 8-fold lower level of CLIP2. Proteins (especially differential proteins of ITGA11, ITGB2, and IL32) showed strong negative correlation with lipids, particularly LPC(O-18:1) and LPC(16:0-SN1). These lipids correlated negatively with 2,3-dihydroxybenzoate. Other highly correlated and discriminant metabolites (e.g., taurodeoxycholate, taurochenodeoxycholate) and lipids (e.g., Cer(18:1;2O/24:1)) were also statistically different.

### Elevated fecal calprotectin associated with increased inflammation-related proteins, SMs; decreased bile acids, amino acids, and TGs

Fecal calprotectin has emerged as a valuable tool in managing IBD, including screening, diagnosis, disease activity and severity assessment, treatment monitoring, and relapse prediction of IBD patients ^6,32^. We explored the molecular perturbations of two IBD subpopulations, i.e., elevated and normal fecal calprotectin levels (**Fig. 3a**). We found 34 differentially expressed proteins between the two groups. Most of them (32 proteins) were up-regulated in patients with elevated fecal calprotectin levels (**Fig. 3b**). These differential proteins belonged to some of the innate and adaptive immune responses, such as TNF, MAPK, IL-17, and NF-κB signaling pathways. Change in proteins involved in the sphingolipid signaling (TNFRSF1A, TRAF2) and PI3K-Akt signaling (CSF1, HGF, OSM, TGFA) was also observed. Besides, 15 differential metabolites were found (**Fig. 3c**). Abundances of chenodeoxycholate and conjugated bile acids (i.e., glycocholate, taurochenodeoxycholate, and glycochenodeoxycholate) decreased (about 2.5-fold), except glycohyodeoxycholate increased slightly in the elevated calprotectin group. Higher levels of two amino acids (creatinine, glutamate) and two DNA metabolism-related metabolites (cytidine, deoxyguanosine-monophosphate) were also observed. Lipids, especially lysoPC, were also found in the metabolomics modality. The lipid modality contributed the most significant number of DAs, with 47 differential lipids (**Fig. 3d**). They presented a clear contrast mostly linked to the low abundance of TG and high abundance of SM. Levels of other lipid subclasses [i.e., PC, PC(O-), LPC, LPE, Cer, and cholesteryl esters (CE)] were also increased.

**Fig. 3.**
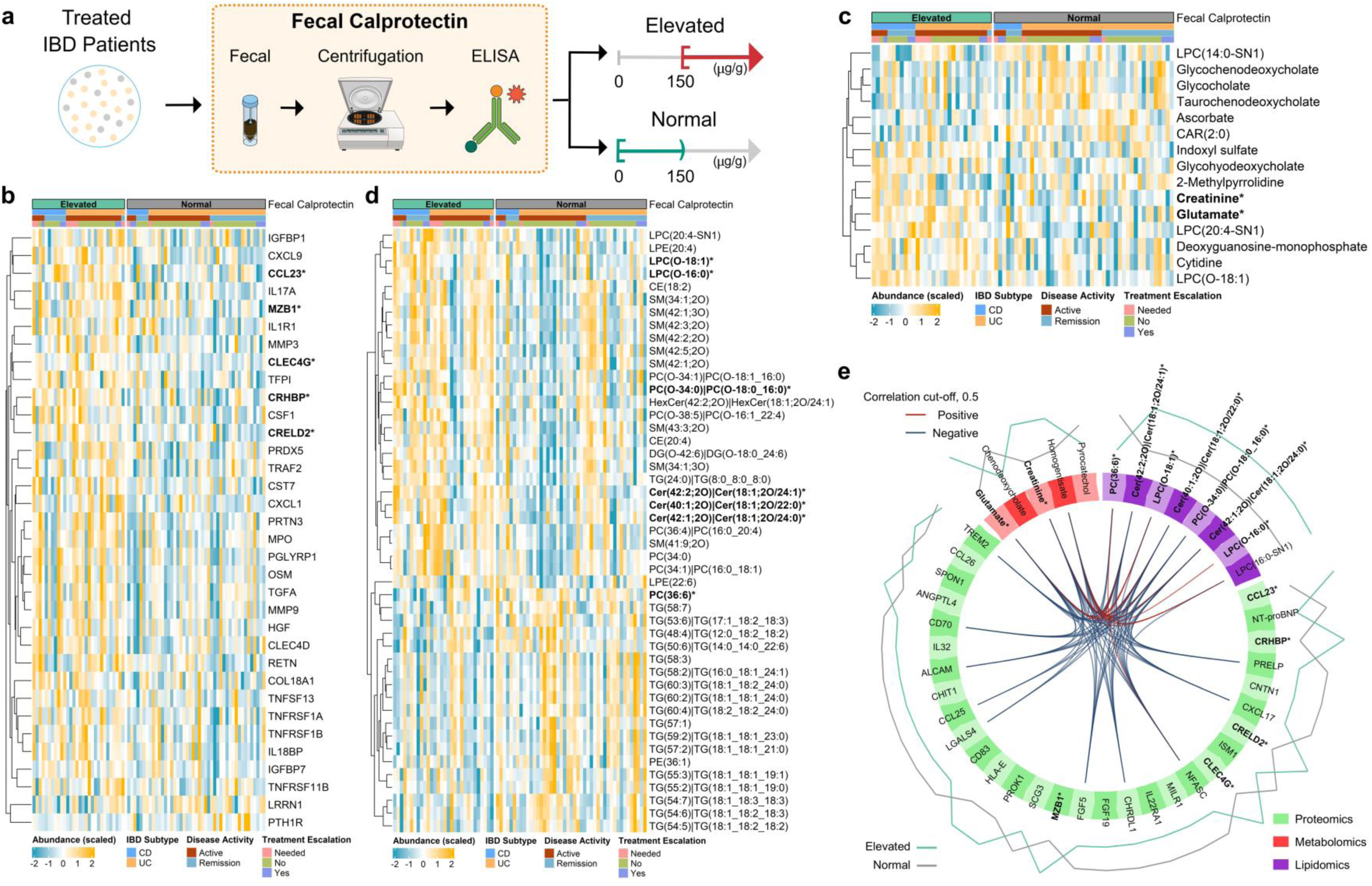
Single and integrative omics analysis presenting molecular perturbances associated with fecal calprotectin levels. **(a)** Sample collection and grouping. Heatmap of differential **(b)** proteins, **(c)** metabolites, and **(d)** lipids. **(e)** DIABLO-selected biosignatures. Abbreviations: IBD: inflammatory bowel disease, UC: ulcerative colitis, CD: Crohn’s disease. * and bold: analytes detected in both single- and multi-omics data analysis.

Integrative omics analysis revealed a good correlation between different data blocks, in which negative correlations were generally observable between proteomics and two other data layers (**Fig. 3e**, **Supplementary Fig. S1b**). However, a strong correlation was the positive correlation of glutamate and creatine with Cer species. Proteins served as the major modality to distinguish 2 groups. Thirty multi-omics-identified proteins were related to immune activation. Only 5 of them were differential analytes. Multi-ome biosignatures also included five metabolites, such as 2 amino acids (creatinine and glutamate; also were differential analytes), potential bacterial-source metabolites (pyrocatechol), and chenodeoxycholate. Eight selected lipids belonged to the Cer, PC, and LPC species, seven of which were differential lipids.

### Active IBD disease state was characterized by an upregulation of inflammation-related proteins, a decrease in SM, and an increase in TG levels

The molecular response underlying patients in active or remission states was investigated (**Fig. 4a**). While lipids showed a significant contribution with 57 differential analytes, 18 and 7 differential proteins and metabolites were observed. Overall, patients with active disease were characterized by the upregulation of all inflammatory-related proteins, a decrease in SM, and an increase in TG levels compared to the remission group (**Fig. 4b-d**). The significantly altered proteins, such as CXCL9, CXCL1, IL17A, TNFSF11, ITGB2, and ADGRE2, were prominently related to activated immune response (**Fig. 4b**). Regarding differential metabolites, muricholate and taurocholate were elevated (2-fold), whereas coprocholate was slightly declined (**Fig. 4c**). Levels of lactate, proline, and acetylcarnitine were decreased. We observed that patients who did not need treatment escalation contributed mainly to the augmented levels of TGs in the active group, whereas abundances of TGs were reduced in the remission group regardless of biologics status (**Fig. 4d**). Lipidomics analysis also revealed a lower abundance of PCs, LPCs, and CEs in the active group.

**Fig. 4.**
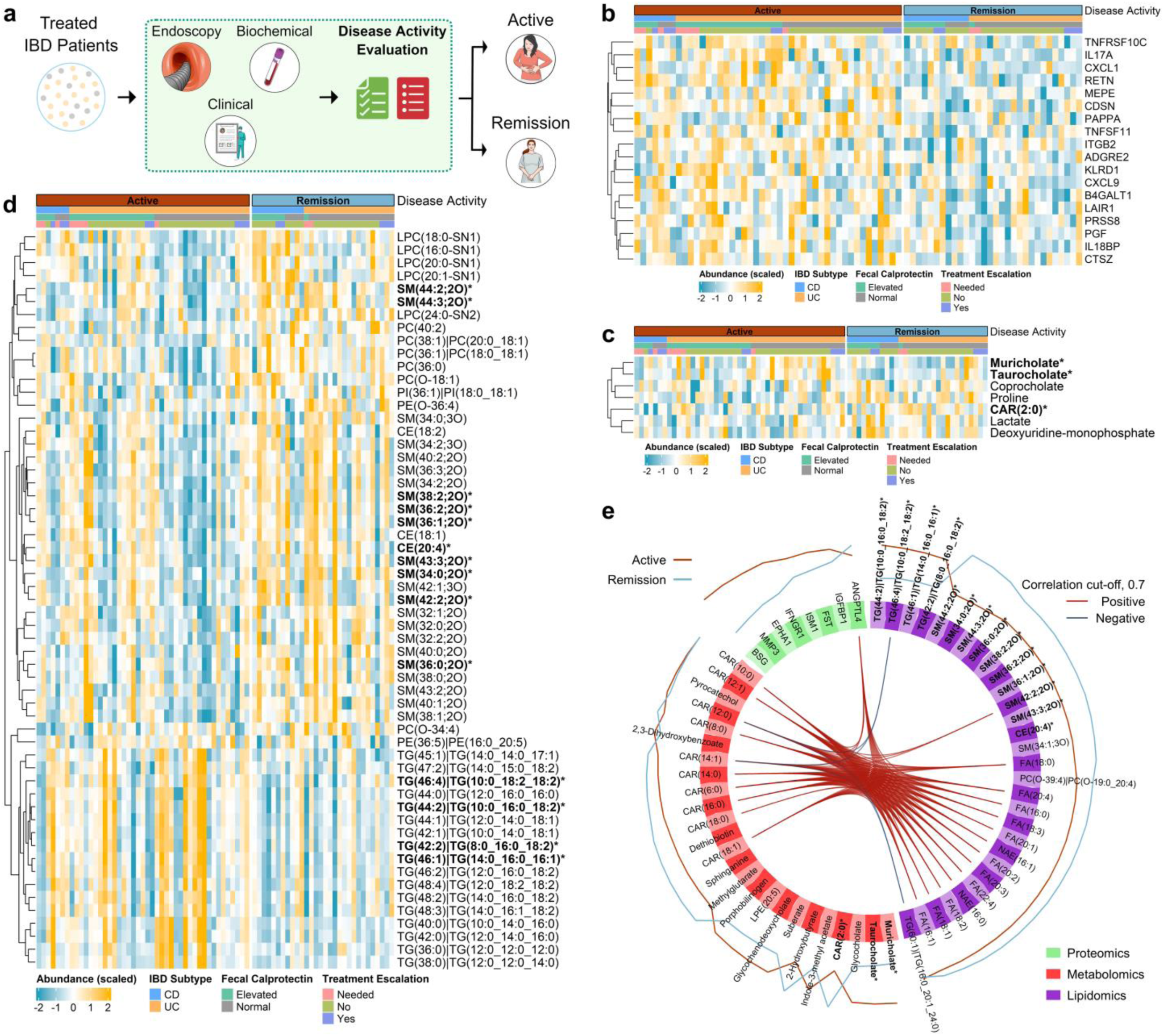
Single and integrative omics analysis presenting molecular perturbances associated with disease activity states. **(a)** Sample collection and grouping. Heatmap of differential **(b)** proteins, **(c)** metabolites, and **(d)** lipids. **(e)** DIABLO-selected biosignatures. Abbreviations: IBD: inflammatory bowel disease, UC: ulcerative colitis, CD: Crohn’s disease. * and bold: analytes detected in both single- and multi-omics data analysis.

In multi-omics analysis, a pair-wise correlation from DIABLO showed a strong correlation between omics layers (**Supplementary Fig. S1c**, **Fig. 4e**). Metabolites and lipids were major drivers for differentiating active and remission disease states. The biosignature comprised 8 proteins, 25 metabolites, and 30 lipids. None of the identified proteins were differentially expressed (**Fig. 4e**). Acylcarnitines were detected primarily within the metabolomics data, yet their abundance between the 2 groups was subtle, except for acetylcarnitine. Two bile acids (muricholate and taurocholate) were also at the top, indicating differences between groups. Within the lipidomics layer, 4 TGs and 9 SMs showed the highest differences between the active and remission groups. Besides, fatty acyls were identified in the multi-omics analysis. They showed a strong positive correlation with ANGPTL4 and acylcarnitines.

### Treatment escalation promoted upregulation of inflammatory response-related proteins, TGs, and decreased amino acids, purine metabolites

We explored molecular perturbations between patients who received treatment escalation with samples collected before and after the therapeutic initiation (**Fig. 5a**). Treatment escalation triggered significant alterations in proteome, metabolome, and lipidome response in IBD patients (**Fig. 5b-d**). We identified 16, 24, and 20 differential proteins, metabolites, and lipids. The majority of altered proteins, which were up-regulated, were linked to the perturbations of the cellular receptor signaling pathway (e.g., COL1A1, FLT3LG, ITGB6, LAMA4), PI3K-Akt signaling pathway, and focal adhesion (**Fig. 5b**). A consistent decrease in levels of differential metabolites was observed (**Fig. 5c**). They were linked to lipid metabolism [e.g., LPC(O-16:0), LPC(O-16:1), LPE(O-18:1), and CAR(12:0;O)], arginine biosynthesis (arginine, aspartate, ornithine, and glutamine), pentose phosphate pathway (hexose-monophosphate, gluconate, and glycerate), purine metabolism (hypoxanthine, glutamine, and deoxyguanosine monophosphate), and amino acid alterations (e.g., creatinine, DOPA, and oxoproline). Lipidome alterations were mainly characterized by a higher level of TGs with a 2-3-fold increase (**Fig. 5d**).

**Fig. 5.**
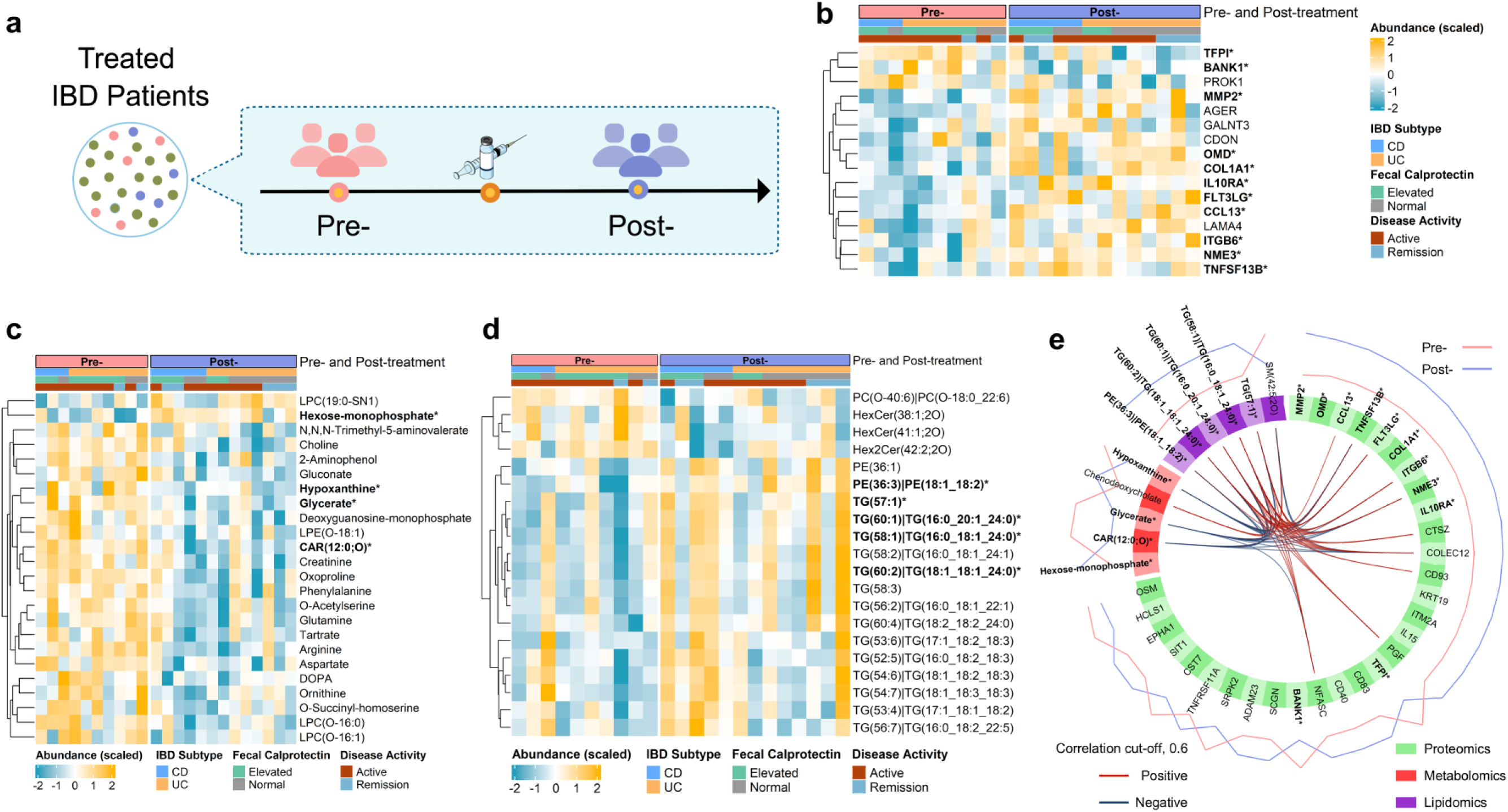
Single and integrative omics analysis presenting molecular perturbances for the pseudo pre- and post-treatment escalation design. **(a)** Sample collection and grouping. Heatmap of differential **(b)** proteins, **(c)** metabolites, and **(d)** lipids. **(e)** DIABLO-selected biosignatures. Abbreviations: IBD: inflammatory bowel disease, UC: ulcerative colitis, CD: Crohn’s disease. * and bold: analytes detected in both single- and multi-omics data analysis.

A high correlation of metabolites and proteins, along with good separation between the two groups were observed, indicated by the distinction in the centroids of each group, yet with an observable moderate amount of overlap between each sample (**Supplementary Fig. S1d**, **Fig. 5e**). Most metabolites (4/5, e.g., hypoxanthine) and lipids (5/6, mainly TGs) in the multi-ome biosignatures were also differential analytes, whereas 11/30 proteins had differential abundance. Lower abundance of hypoxanthine and glycate were correlated strongly with decreased abundances of TGs and increased levels of proteins such as CCL13, ITGB6, and COLEC12.

### Patients who needed to escalate their treatment showed downregulation of signaling receptor binding proteins, lower levels of TGs, higher levels of SMs and PCs

The multi-ome profiles of patients who needed treatment escalation and those who did not were compared (**Fig. 6a**). The adjusted linear model revealed 25 proteins, mostly linked to signaling receptor binding, differentially expressed (11 up- and 14 down-regulated) between two groups (**Fig. 6b**). Proteins linked to cytokine-cytokine receptor interaction were down-regulated (2 to 10-fold; CXCL6, CD40LG, CCL13, TNFRSF14, and TNFSF13B) in patients who needed treatment escalation, except CXCL9 increased 6-fold. Four proteins involved in MAPK and PI3K-Akt signaling pathways (i.e., FLT3LG, EGF, EGFR, and PDGFA) were also consistently down-regulated. Changes in the proteins mentioned above might also indicate the alteration of NF-κB and JAK-STAT signaling pathways. For metabolomics data, we determined 10 differential metabolites (**Fig. 6c**). A marginal increase in the abundance of three amino acids (DOPA, arginine, and ornithine) was observed, which might imply a change in arginine metabolism. Regarding lipidome alteration, 31 lipids were statistically significant (**Fig. 6d**). TG species showed a lower level (2-3-fold) in patients who required treatment escalation, whereas other lipid subclasses, such as PC(O-), PC, and Cer had increased abundance.

**Fig. 6.**
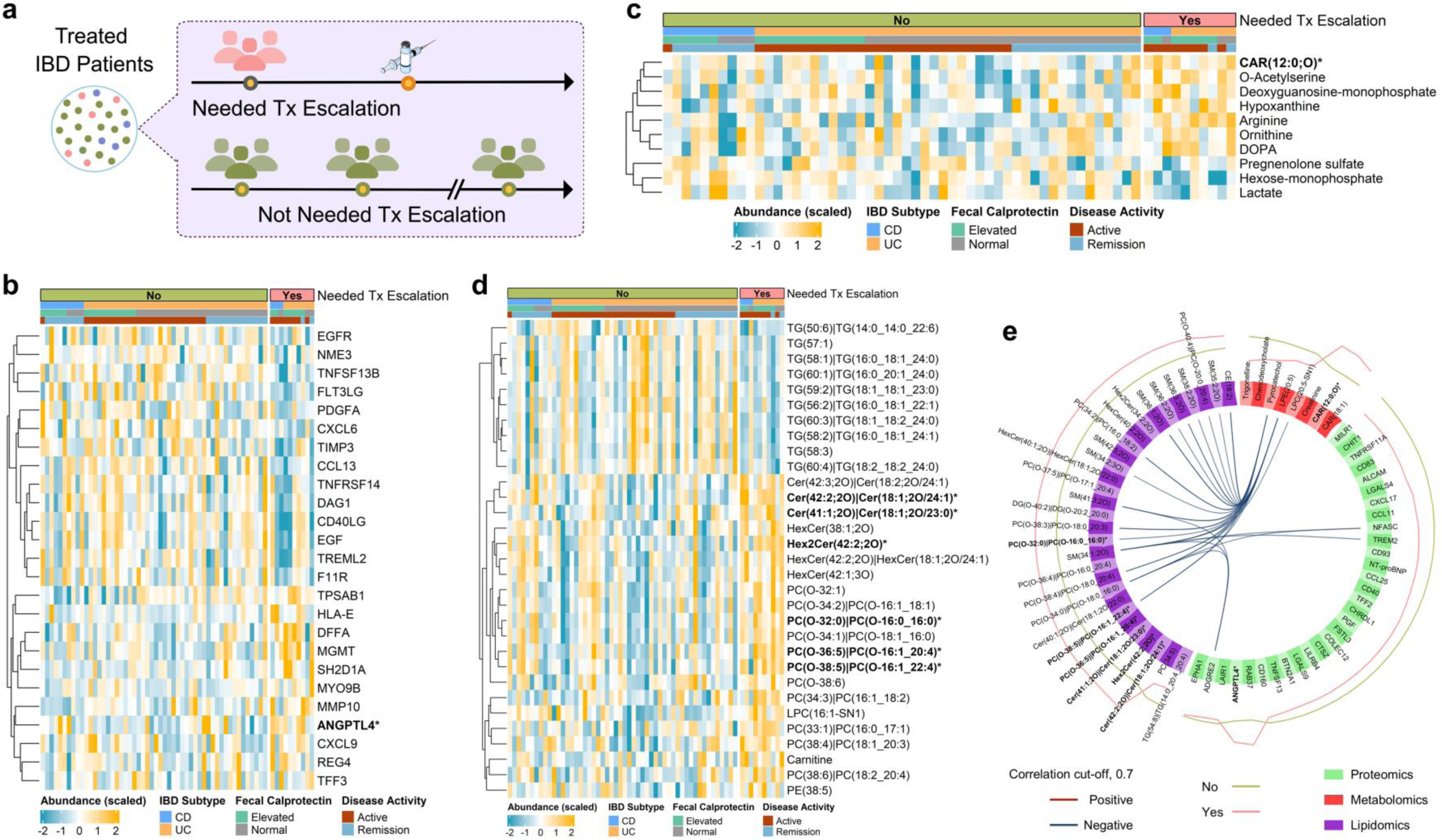
Single and integrative omics analysis presenting molecular perturbances between IBD patients who needed treatment escalation and those who did not. **(a)** Sample collection and grouping. Heatmap of differential **(b)** proteins, **(c)** metabolites, and **(d)** lipids. **(e)** DIABLO-selected biosignatures. Abbreviations: IBD: inflammatory bowel diseases, Tx escalation: treatment escalation, UC: ulcerative colitis, CD: Crohn’s disease. * and bold: analytes detected in both single- and multi-omics data analysis.

A good correlation between proteome, metabolome, and lipidome profiles was observed within two groups (**Supplementary Fig. S1e**, **Fig. 6e**). The negative correlation pattern was mainly observed, such as between NFASC and TREM2 with SM(34:1;2O). Proteins identified by multi-omics analysis, which mainly down-regulated, were linked to the immune response. While metabolites range from acylcarnitines to bile acids (chenodeoxycholate) and bacterial-sourced metabolites (pyrocatechol), selected lipids were mainly SM and PC(O-) species, whose higher abundance in patients needed treatment escalation.

### Integrative biosignatures captured differences between groups of five scenarios

The predictive performance of selected multi-ome biosignatures is shown in **Table 2**. Differences between CD and UC groups, as well as between individuals with elevated and normal fecal calprotectin levels, could be captured by the selected multi-ome biosignatures (AUC = 0.80, P-value = 0.008; AUC = 0.73, P-value = 0.03; respectively). However, the model failed to differentiate active from remission disease state groups (AUC = 0.56, P-value = 0.42). Regarding treatment escalation, the biosignatures exhibited an excellent performance in distinguishing post-from pre-treatment escalation groups (AUC = 0.90, P-value = 0.01). Additionally, a good capability was observed in distinguishing patients needing to escalate their treatment from those not (AUC = 0.77, P-value = 0.02). Among these five integrative biosignatures, 8 analytes were detected in at least 3 of 5 scenarios (**Supplementary Fig. S2**). They included NFASC, CD83, chenodeoxycholate, ANGPTL4, pyrocatechol, Cer(18:1;2O/24:1), ISM1, and EPHA1. Their abundances across 5 scenarios are shown in **Supplementary Figs. S3-10**.

**Table 2.** Performance regarding the classification of the biosignatures in five scenarios.

|  | **AUC<sup>1</sup>** | **P-value** |
| --- | --- | --- |
| **IBD subtypes, CD *vs.* UC** | 0.80 | 0.008 |
| **Fecal calprotectin levels, elevated *vs.* normal** | 0.73 | 0.03 |
| **Disease state, active *vs.* remission** | 0.56 | 0.42 |
| **Post- *vs.* pre-treatment escalation** | 0.90 | 0.01 |
| **Needed *vs.* not needed treatment escalation** | 0.77 | 0.02 |
| <sup>1</sup> Average AUC via cross-validation procedure. |  |  |

## DISCUSSION

Owing to the heterogeneous nature of IBD, the current disease classification could preclude the endeavors to understand the pathophysiology of the disease. A classification comprising multiple biomarkers that reflect the IBD spectrum holds promise to account for diverse manifestations of IBD ^5^. Hence, using multi-ome molecular profiles may be a better approach. Our study set out to investigate five commonly seen clinical scenarios and provide the evidence about the molecular differences of the comparable groups at the functional multi-ome data. It pinpointed the serum multi-ome profiles underlying the disease and associated with either the fecal calprotectin levels or disease activity states, as well as molecular profiles upon treatment escalation.

We first aimed to elucidate the molecular differences between UC and CD. Barrier dysfunction is known to be the primary mechanism underlying IBD pathophysiology ^4^. In UC, the injury only extends to the mucosal layer, whereas transmural inflammation occurs in CD ^12^. In our study, cell adhesion molecules were consistently higher in UC compared to CD, suggesting the importance and potential utility of these proteins. Among them, the increased expression of ITGA11, ITGB2, and IL32 demonstrated strong negative correlation with LPC(O-18:1) and LPC(16:0-SN1). The alterations of PC species and their derivatives suggests the disruption of the gut mucus layer. Previous studies found that PCs and LPCs species decreased in UC versus CD, whereas we observed a consistent decrease of LPC species and PCs showed species-dependent patterns ^16^. The shift of phospholipid composition in intestinal mucus could lead to an increase in the permeability of the intestinal mucus, promoting mucosal inflammation ^33^. Bile acids are mainly re-absorbed via the ileum, a well-known location for CD compared to UC. In our study, changes in bile acid homeostasis indicated by altered levels of bile acids and FGF19 expression between two disease subtypes also might induce chronic inflammation, resulting in an impaired mucosal barrier ^34,35^.

Fecal calprotectin could be used as a surrogate endpoint presenting gastrointestinal tract inflammation in IBD. Mechanistically, non-apoptotic epithelial cell death, oxidative stress, and homeostasis pathway dysregulation result in an excessive release of calprotectin, promoting inflammation ^36^. Establishing the association between serum multi-ome profiles and fecal calprotectin to shed light on the systemic molecular alterations upon inflammation. In our study, patients with elevated fecal calprotectin presented up-regulation of proteins involved in inflammation-promoting signaling pathways. Inflammation and microbiome dysbiosis might lead to malabsorption and disrupt bile acid function in the gastrointestinal tract. This might result in systemic changes in amino acids and DNA-related metabolites, as observed in our study ^14,15^. Besides, determining the relationship between serum multi-omic profiles and disease activity could illuminate systemic molecular alterations associated with active disease manifestations. We found that active disease was characterized by up-regulation of Th1 and Th17 immune activation-related proteins such as CXCL9, CXCL1, ITGB2, and IL17A. Between these two scenarios, we observed differences, yet significance in the direction of change of TGs and SMs. In the gut, the immunomodulatory role of SMs, either of host- or microbiome-origin, has been recognized as contributing to cell growth, differentiation, apoptosis, and membrane structure integrity ^16,37,38^. Additionally, intestinal spontaneous inflammations could result from reduced bacterial sphingolipids. IBD patients suffer from lipid abnormalities that correlate with disease activity ^39^. We observed consistently increased levels of TGs in the active disease state, which may be helpful biomarkers and associated with the pathophysiology of the active disease state.

We also designed the pre- and post-treatment scenarios to investigate the molecular perturbations upon treatment escalation. Treatment escalation appeared to be associated with an increase of inflammatory response-related proteins and TGs; and a decrease of amino acids and purine metabolites. The high abundance of TGs after treatment escalation could be linked to the reduced abundance of TGs in the elevated fecal calprotectin group as a response to inflammation alleviation. Moreover, needed-treatment-escalation patients had a lower level of TGs compared to patients who did not require treatment escalation. It should also be noted that increased TGs in the active-disease-state patients were attributed predominantly to individuals who did not need treatment escalation. Therefore, TGs might have the potential for treatment monitoring and complication detection. However, a more sophisticated analysis in the context of improved study design and sample size is warranted to establish the association ^40^. Patients required to escalate their treatment tend to suffer more severity of disease and thus need to escalate their treatment. Nevertheless, disease severity assessment is a hard-to-implement process requiring multiple criteria ^32^. Therefore, fast and convenient biosignatures might open the door for early treatment intervention. We observed the decreased levels of signaling receptor binding proteins and higher levels of SMs and PCs. However, CXCL9 increased 6-fold in needed-treatment-escalation patients. The abundance of this protein was also significantly increased in patients with active disease states and elevated calprotectin. Notably, CXCL9 significantly increased the risk of disease development in CD patients ^41^. Therefore, pre-clinical to clinical activation of CXCL9 might be of potential for clinical utilization.

Integrative biosignature holds the potential to account for diverse manifestations and capture the holistic change of IBD. We evaluated predictive performance to give an idea about the potential of the proposed biosignature. This *priori* information might be of potential use for further studies.

Some common analytes detected might be potential candidates for IBD management. For example, neurofascin (NFASC), which was only detected by integrative analysis, was a biomarker candidate among 3 of 5 investigated scenarios with consistently lower levels in elevated-calprotectin, active-disease-state, and needed-treatment-escalation patients. Also, the NFASC levels were lower in CD patients and tended to be normalized in the post-treatment escalation group. Neurofascin is known for its mechanistic and clinical role in peripheral neuropathy, which is one of the most common yet unrecognized neurological disorders in the IBD population, especially in CD patients ^42–46^. A previous study showed that aged monkey brain tissues presented increased interaction between neurofascin and mutant α-synuclein, and transfected neurofascin could ameliorate its neuritic toxicity in neuronal cells. Alpha-synuclein is known to cause Parkinson’s disease, which is co-occurrence and increased risk in IBD patients ^47^. Additionally, early anti-inflammatory treatment, especially anti-TNF-α agents, has been associated with decreased Parkinson’s disease risk in the IBD population ^48,49^. Another protein, ANGPTL4, known for its regulatory role in inflammatory responses across various diseases ^50^, was augmented in patients with elevated calprotectin and in patients who needed treatment escaltion, yet decreased in active cases. Studies showed elevated plasma levels of ANGPTL4 in mild/moderate IBD patients, returning to normal levels in severe cases ^51^. Increased ANGPTL4 expression in the inflamed colons correlated with the reduced infiltration of CD8+ cells, suggesting its potential role in modulating gut immune responses ^51^.

Some limitations in our study should be noted. First, the genetic background of the patients was unavailable. This hampered the exploration of the genetics and molecular profiles. Second, the limited number of participants might restrict the generalizability of some findings and make it hard to detect subtle but potentially important differences in the multi-ome profiles. Besides, the inclusion of treated patients due to the convenient sampling setting prevented us from exploring the effects of the underlying disease itself and the effects of the treatment. Although the molecular profiles reported in our study were comprehensive, the cross-sectional study design made it necessary to improve the study design to draw the causative relationship.

## CONCLUSION

This study conducted a functional multi-omics approach toward deep molecular phenotyping of IBD. The comprehensive analysis revealed distinct molecular biosignatures across five typical IBD scenarios. Our findings provide deeper insights into serum multi-ome profiles underlying the disease and in association with either the fecal calprotectin levels or disease activity states, and molecular profiles upon treatment escalation. These biosignatures pointed towards disturbances in key pathophysiological processes like immune response, bile acid homeostasis, amino acids, and lipids alteration, potentially underlying the heterogeneous spectrum of IBD. Some potential candidates, such as NFASC, ANGPTL4, chenodeoxycholate, and TGs hold promise as potential candidates for IBD management. Future validation in more extensive and longitudinal cohorts and translation into clinical tools can pave the way for a more precise approach to IBD diagnosis, stratification, and treatment.

## ACKNOWLEDGEMENTS

This work was supported by Eisai Korea Inc. The funder was not involved in the study design, data acquisition, data analysis, data interpretation, manuscript writing and editing, or publication.

The biospecimens and data used in this study were provided by the Biobank of Inje University Busan Paik Hospital, member of Korea Biobank Network (KBN4_A08).

## AUTHOR CONTRIBUTIONS STATEMENT

**N.T.N.T.**: Data curation, Methodology, Investigation, Formal analysis, Validation, Writing – Original Draft, Writing – Review & Editing. **E.J.C.**: Data curation, Methodology, Investigation, Formal analysis, Visualization, Writing – Review & Editing. **N.Q.T.**: Data curation, Methodology, Software, Validation, Writing – Review & Editing. **S.J.Y.**: Data curation, Investigation, Validation, Writing – Review & Editing. **D.N.N.**: Methodology, Supervision, Validation, Writing – Review & Editing. **D.H.K.**: Conceptualization, Investigation, Supervision, Resources, Writing – Review & Editing, Funding acquisition. **N.P.L.**: Conceptualization, Methodology, Investigation, Formal analysis, Validation, Supervision, Resources, Writing – Original draft, Writing – Review & Editing, Funding acquisition. **H.S.L.**: Conceptualization, Methodology, Investigation, Formal analysis, Validation, Resources, Supervision, Writing – Original draft, Writing – Review & Editing, Funding acquisition.

## DATA AVAILABILITY STATEMENT

The data supporting this study’s findings are available upon reasonable request.

## SUPPLEMENTARY INFORMATION

Supplementary materials for this article can be found online.

