## Supplementary Material for "Multi-omics phenotyping characterizes molecular divergence underlying different clinical scenarios of inflammatory bowel disease"

### SUPPLEMENTARY METHODS

#### Sample preparation for metabolomics and lipidomics

Metabolomics and lipidomics experiments followed our previously established procedure [22-24]. For metabolite extraction, 50 μL of the serum sample (−80 °C) was first thawed on ice for about 30 min and then vortexed extensively for 10 s. Afterward, 150 μL precooled extraction solvent, which was the pre-dissolved of the 6 internal standards in MeOH, was pipetted into each sample. The mixtures were vortexed thoroughly for 30 s and then centrifuged at 14,000 *rcf* and 4 °C for 2 min. Next, 150 μL of supernatant was taken and evaporated using a nitrogen gas flow. The dried samples were resuspended in 200 μL of 50% MeOH and then vortexed vigorously. The samples were centrifuged at 14,000 *rcf* and 4 °C for 2 min before obtaining 150 μL of supernatant and 20 μL of supernatant for pooled quality control (QC) sample to inject into the LC-MS.

In lipid extraction, pre-aliquoted serum samples (50 μL at −80 °C) were initially thawed on ice for approximately 30 min. Subsequently, 10 μL of a pre-mixed IS of SPLASH and Deuterated Ceramide LIPIDOMIX (Alabama, USA) was spiked to each sample, followed by brief vortexing. After a 20-min incubation on ice and underwent vortexing, 300 μL of methanol (−20 °C) and 1000 μL of MTBE (−20 °C) were put in. The mixtures were vortexed for 10 s and incubated in a shaker for 20 min at 1500 *rpm* and 4 °C. Subsequently, 250 μL of water was added, vortexed for 20 s, and incubated for 10 min on ice. The samples were centrifuged for 2 min at 4 °C and 14,000 *rcf*. Next, 500 μL of supernatants were collected from each sample and dried under a nitrogen gas flow. Methanol/toluene (9:1, *v/v*) of 200 μL was used to resuspend the dried lipid extracts. The samples were then introduced into a centrifuge for 2 min at 4 °C and 14,000 *rcf* to obtain 150 μL of supernatant for instrumental analysis. Additionally, post-extraction pooled quality control (QC) samples were created by mixing and vortexing 20 μL of supernatant from each sample.

**Metabolomics and lipidomics instrumental analysis and data acquisition**

The instrumental analysis consisting of the Shimadzu Nexera LC system (Kyoto, Japan) and the X500R Quadrupole Time-of-Flight mass spectrometer combined with Turbo V™ ion source and TwinSpray probe (SCIEX, MA, USA) was utilized [22-24]. For metabolomics, ACQUITY UPLC HSS T3 column (50 mm × 2.1 mm; 1.8 µm) coupled with ACQUITY UPLC HSS T3 VanGuard pre-column (5 mm × 2.1 mm; 1.8 µm) was used. Mobile phase A of 100% water with 0.2% FA and mobile phase B of 100% ACN with 0.1% FA were implemented [25]. The injection volume for positive (POS) and negative (NEG) ion modes was 3 µL. For lipidomics, the stationary phase of ACQUITY UPLC BEH C18 column (50 mm × 2.1 mm; 1.7 µm) connected to an ACQUITY UPLC BEH C18 VanGuard pre-column (5 mm × 2.1 mm; 1.7 µm) was used. Mobile phase A was ACN/water (60:40, *v/v*) with 10 mM ammonium formate and 0.1% FA, and mobile phase B was IPA/ACN (90:10, *v/v*) containing 10 mM ammonium formate and 0.1% FA [25,26]. Within the POS, 1 µL of the samples were injected. The injection volume for the NEG was 2 µL. Mass calibration was performed after every five injections during the analysis, and data was acquired via data-dependent mode.

### SUPPLEMENTARY FIGURES

**Fig. S1. Diagnostic plots of DIABLO-selected biosignatures in five investigated scenarios. (a)** IBD subtypes, **(b)** fecal calprotectin, **(c)** disease activity state, **(d)** pre- and post-treatment escalation, and **(e)** need treatment escalation. Three left panels show pair-wise correlation coefficients. Three right panels present samples colored by groups with their 95% confidence ellipse.

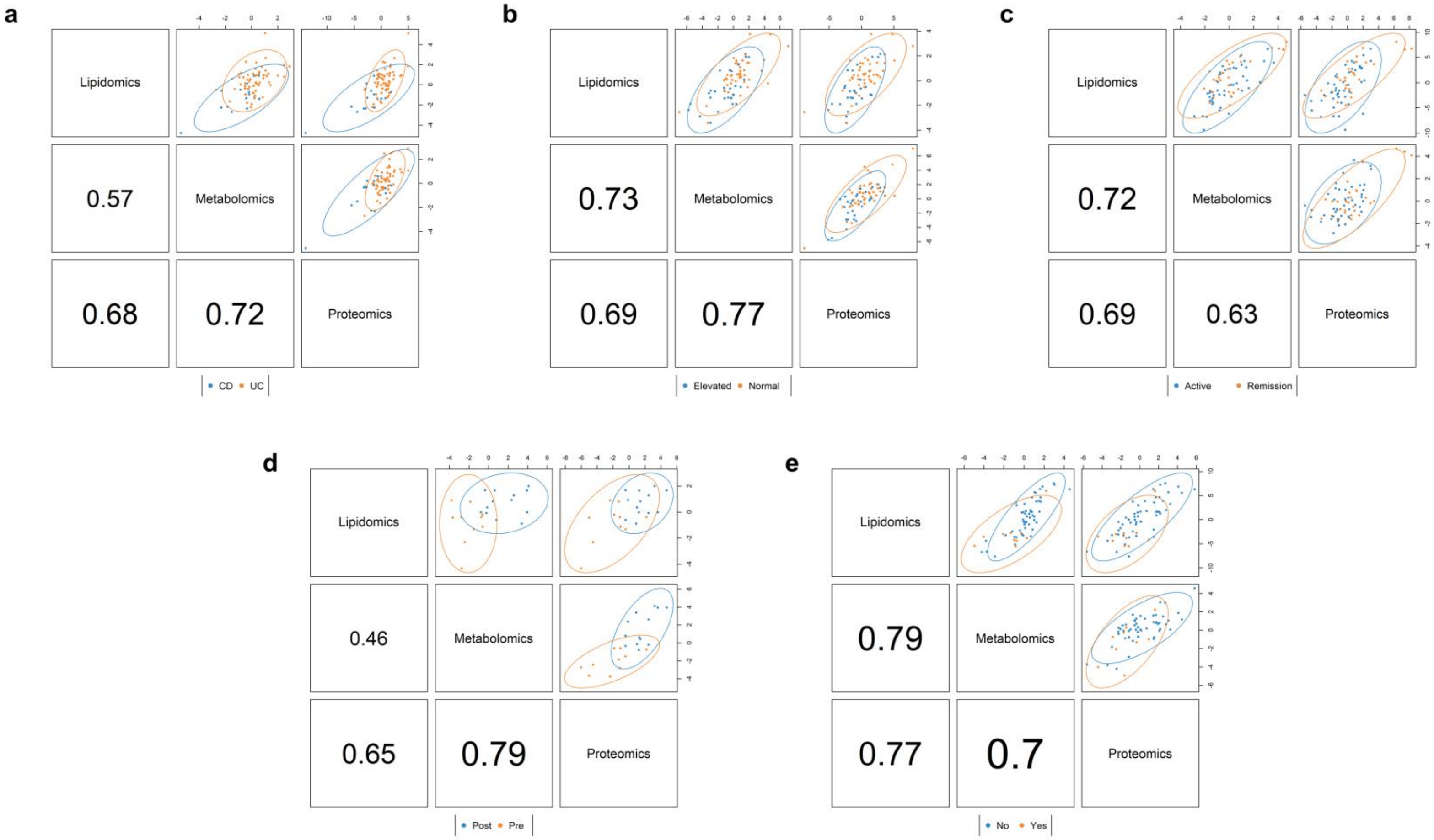

**Fig. S2. Upset plot of DIABLO-selected biosignatures in five investigated scenarios.**

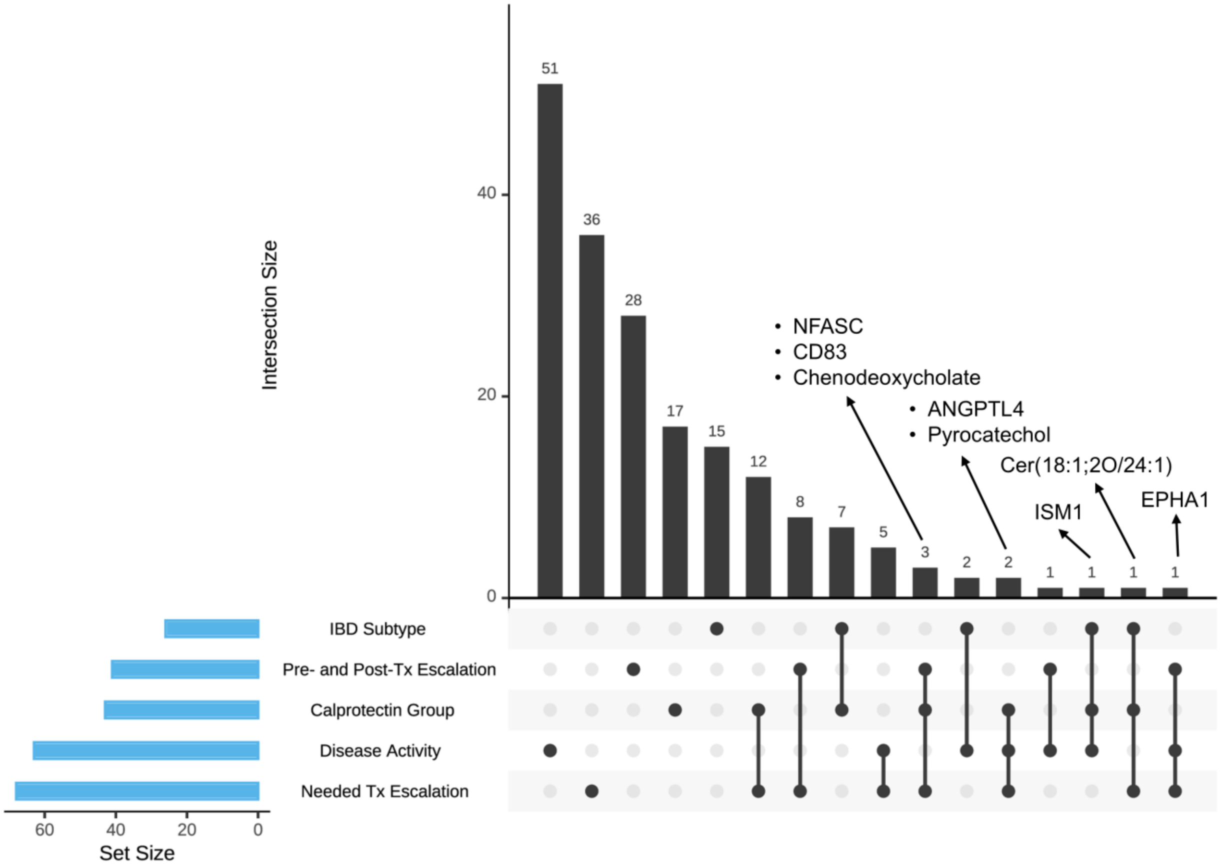

**Fig. S3. Abundances of neurofascin (NFASC) in five investigated scenarios of (a)** IBD subtypes, **(b)** fecal calprotectin, **(c)** disease activity state, **(d)** pre- and post-treatment escalation, and **(e)** need treatment escalation.

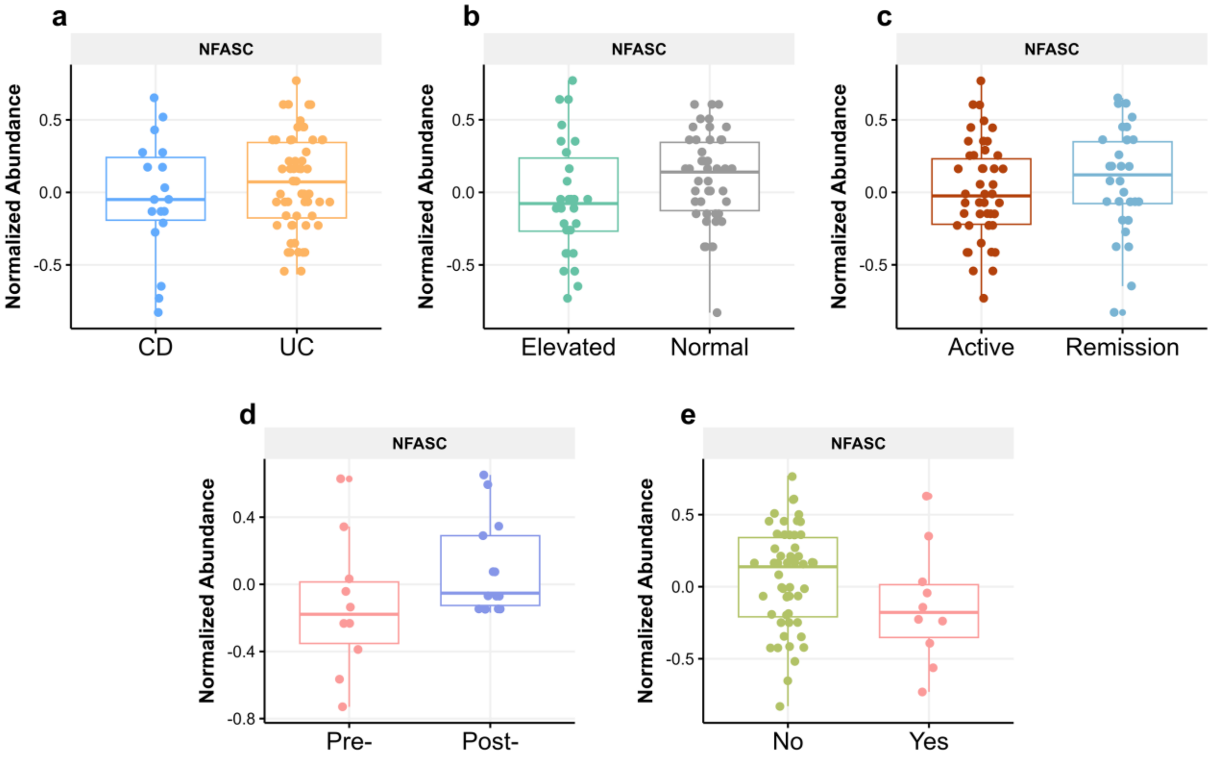

**Fig. S4. Abundances of CD83 antigen (CD83) in five investigated scenarios of (a)** IBD subtypes, **(b)** fecal calprotectin, **(c)** disease activity state, **(d)** pre- and post-treatment escalation, and **(e)** need treatment escalation.

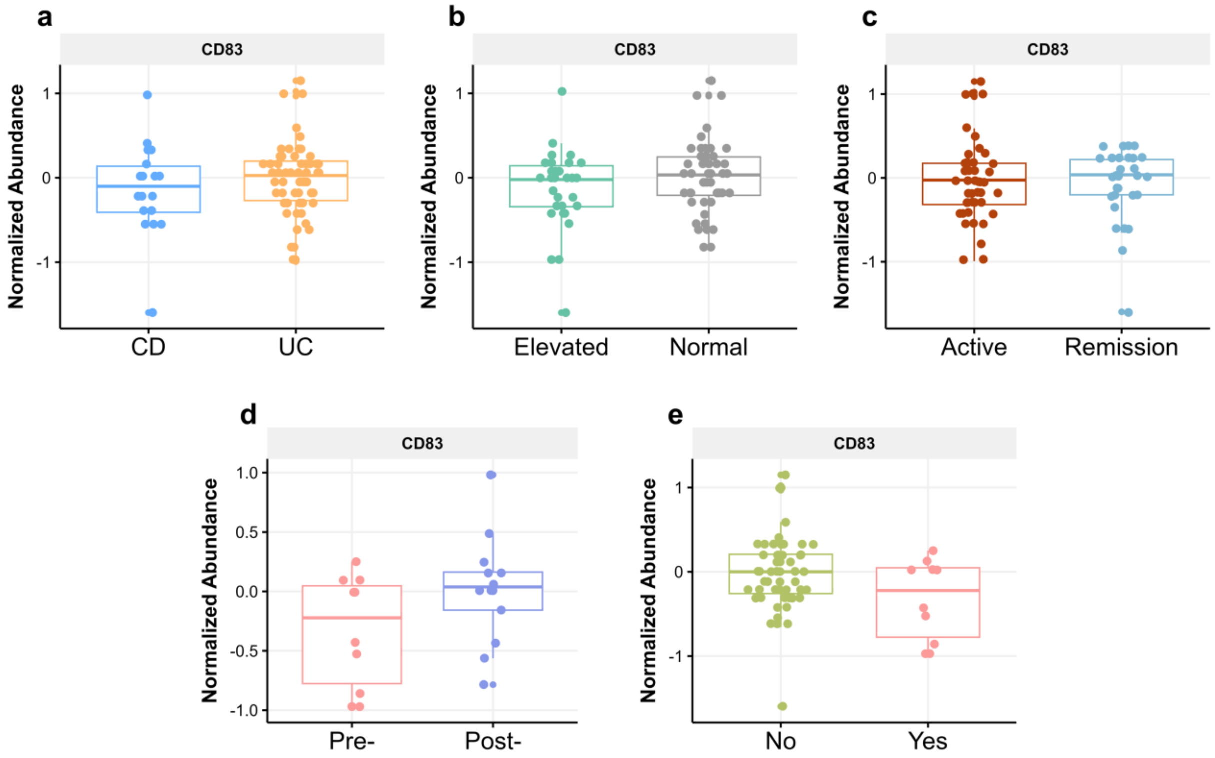

**Fig. S5. Abundances of chenodeoxycholate in five investigated scenarios of (a)** IBD subtypes, **(b)** fecal calprotectin, **(c)** disease activity state, **(d)** pre- and post-treatment escalation, and **(e)** need treatment escalation.

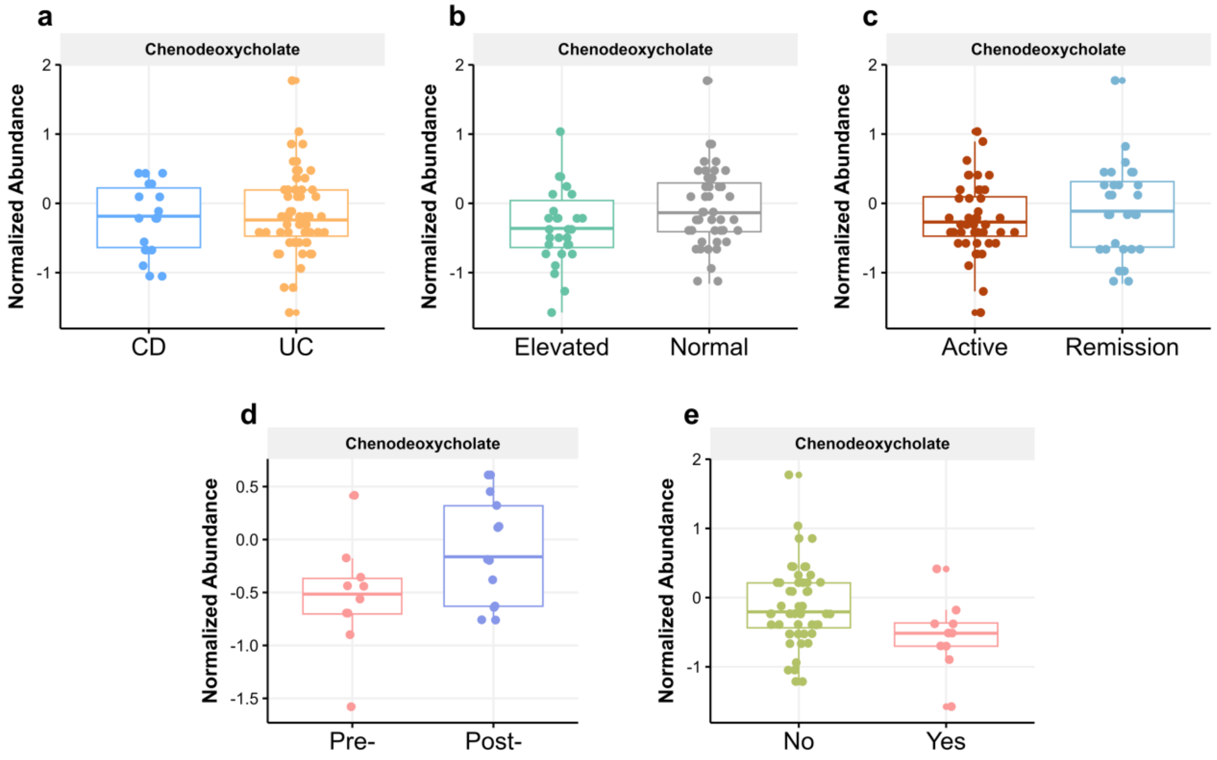

**Fig. S6. Abundances of angiopoietin-related protein 4 (ANGPTL4) in five investigated scenarios of (a)** IBD subtypes, **(b)** fecal calprotectin, **(c)** disease activity state, **(d)** pre- and post-treatment escalation, and **(e)** need treatment escalation.

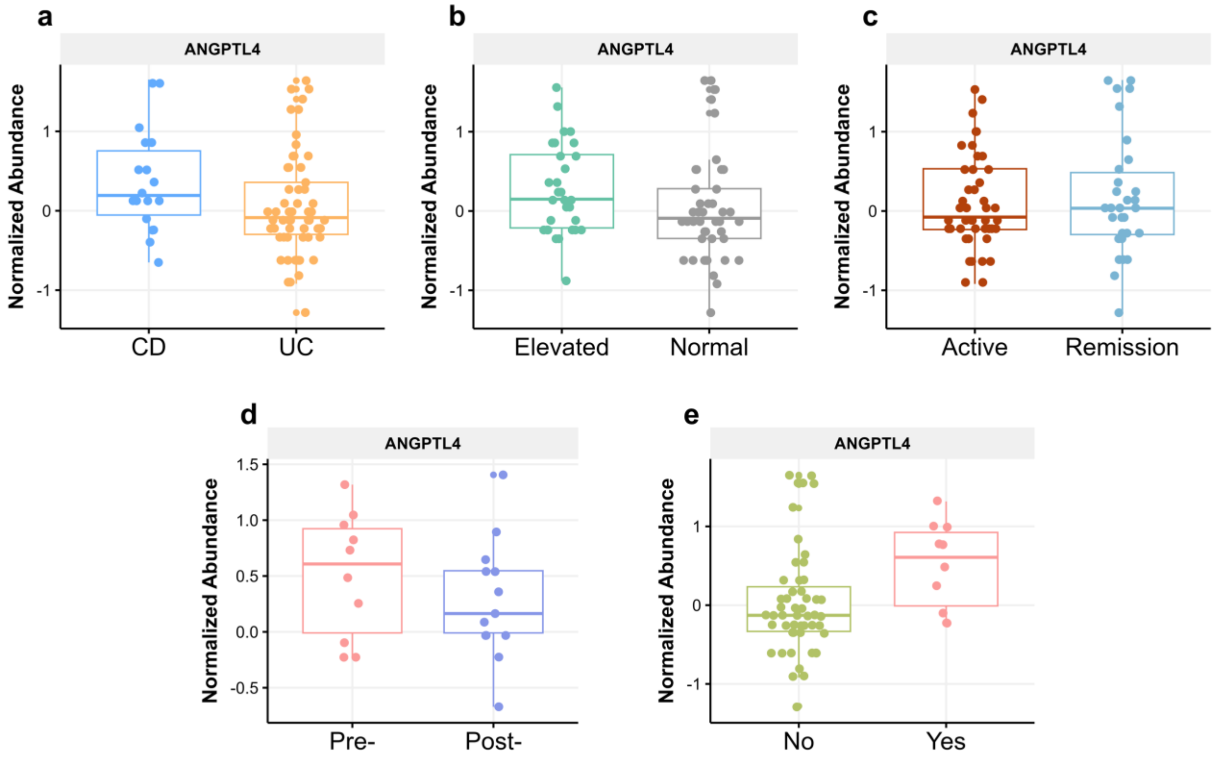

**Fig. S7. Abundances of pyrocatecholin in five investigated scenarios of (a)** IBD subtypes, **(b)** fecal calprotectin, **(c)** disease activity state, **(d)** pre- and post-treatment escalation, and **(e)** need treatment escalation.

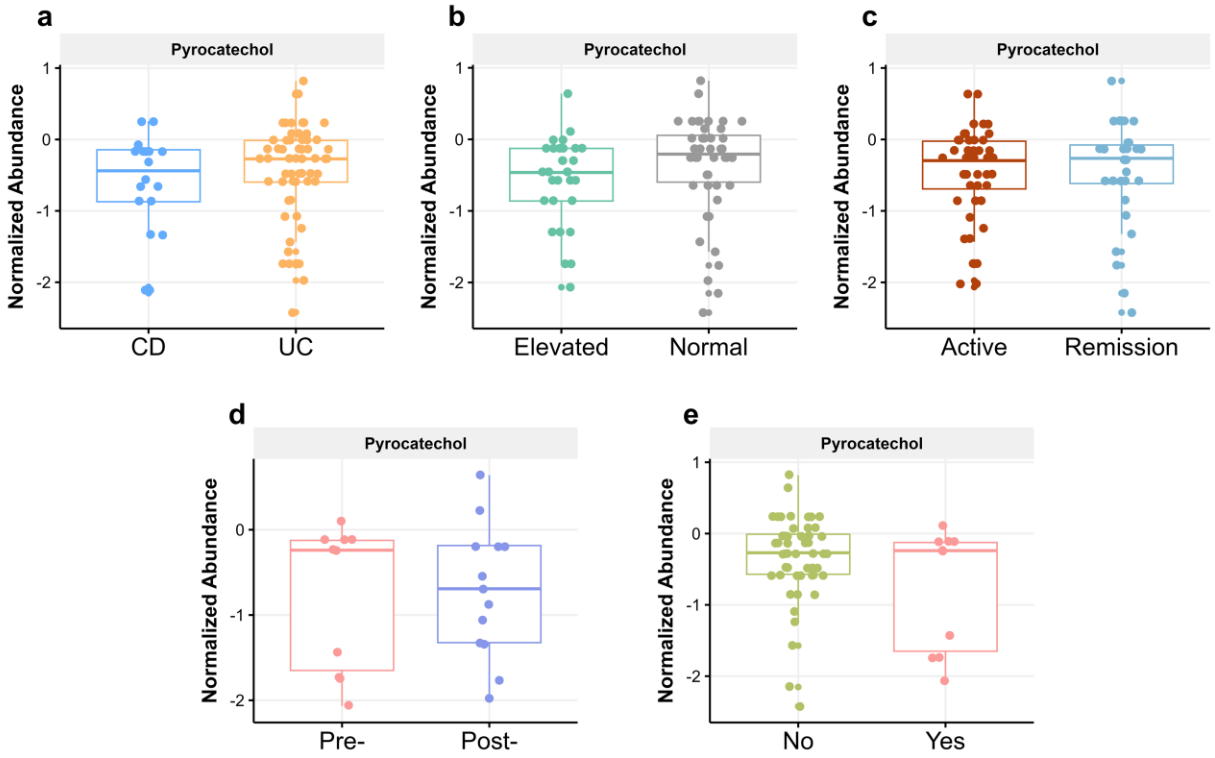

**Fig. S8. Abundances of isthmin-1 (ISM1) in five investigated scenarios of (a)** IBD subtypes, **(b)** fecal calprotectin, **(c)** disease activity state, **(d)** pre- and post-treatment escalation, and **(e)** need treatment escalation.

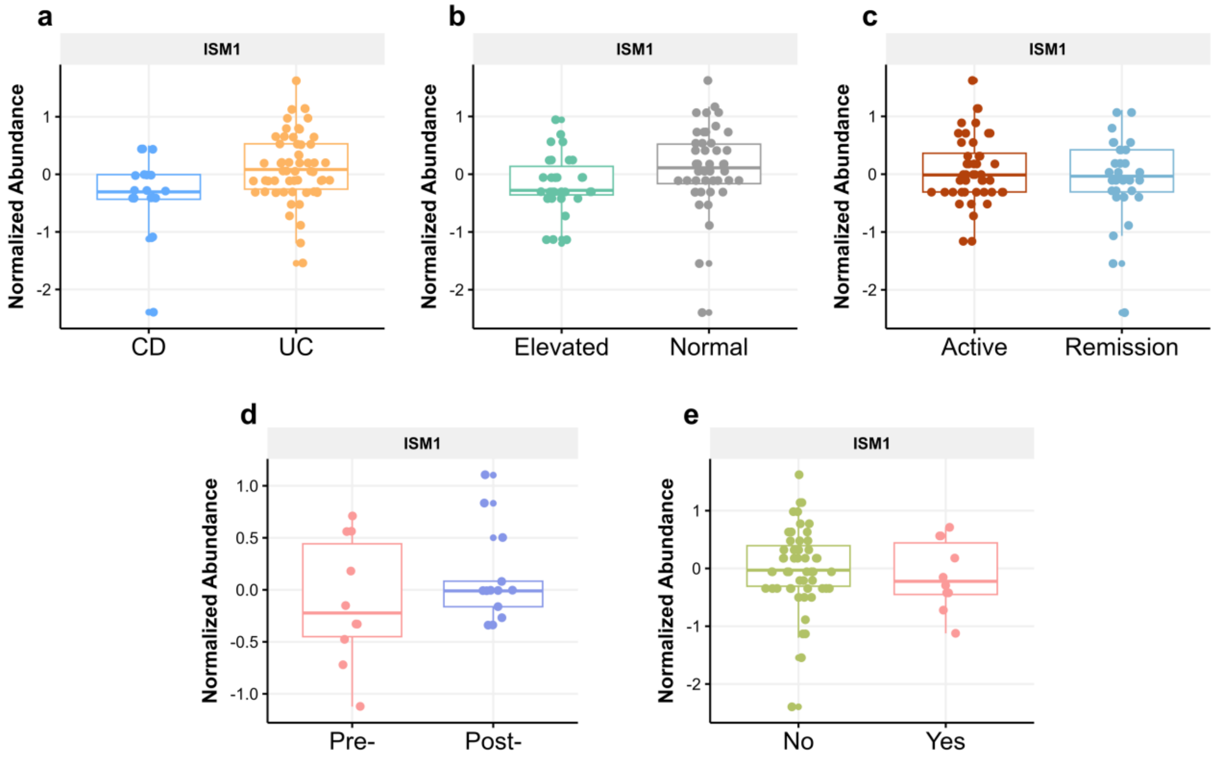

**Fig. S9. Abundances of Cer(18:1;2O/24:1) in five investigated scenarios of (a)** IBD subtypes, **(b)** fecal calprotectin, **(c)** disease activity state, **(d)** pre- and post-treatment escalation, and **(e)** need treatment escalation.

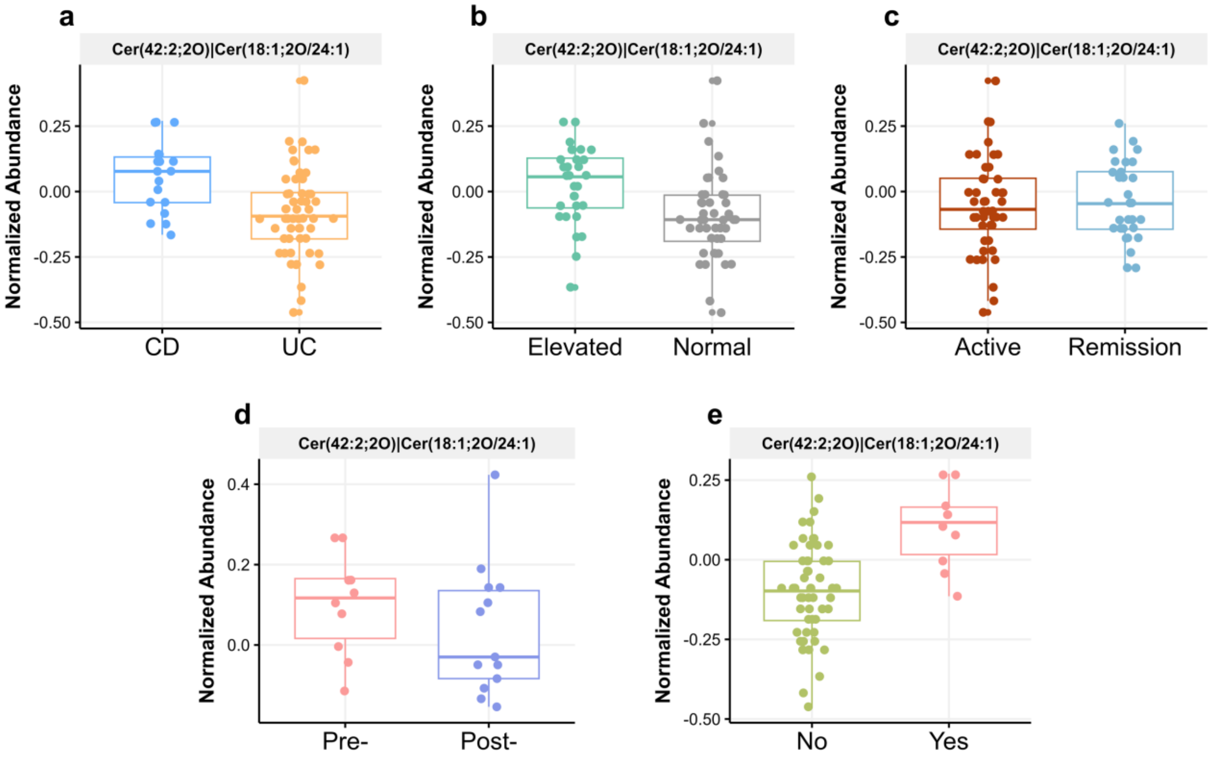

**Fig. S10. Abundances of ephrin type-A receptor 1 (EPHA1) in five investigated scenarios of (a)** IBD subtypes, **(b)** fecal calprotectin, **(c)** disease activity state, **(d)** pre- and post-treatment escalation, and **(e)** need treatment escalation.

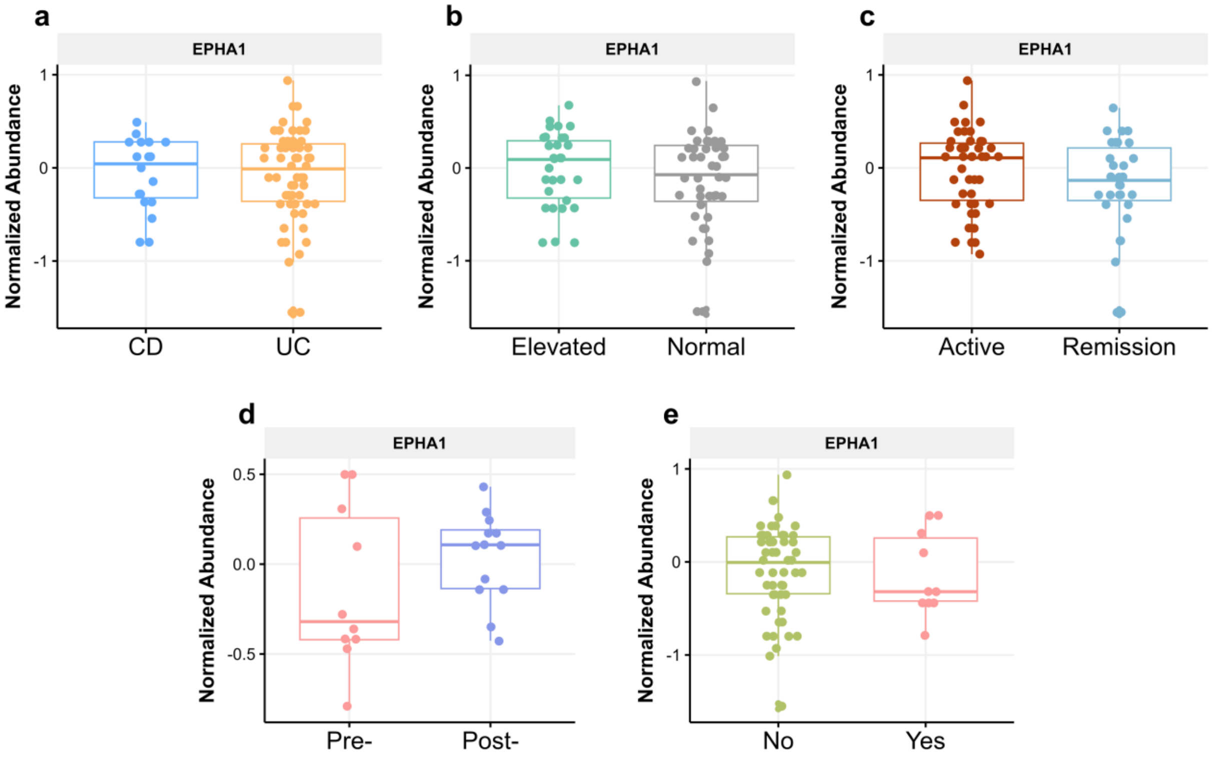

### SUPPLEMENTARY TABLES

**Table S1. Clinical characteristics by fecal calprotectin group.**

| | Elevated, N = 30[1,2] | Normal, N = 45[1,2] |
|---|---|---|
| **Age at recruitment [years]** | 33 (25, 51) | 43 (27, 55) |
| **BMI [kg.m$^{-2}$]** | 22.8 (20.8, 25.6) | 22.0 (20.7, 23.7) |
| **Sex, females** | 7 (23%) | 20 (44%) |
| **Time since diagnosis [years]** | 1.06 (0.22, 2.19) | 1.70 (0.50, 2.75) |
| **Disease subtype** | | |
| CD | 11 (37%) | 7 (16%) |
| UC | 19 (63%) | 38 (84%) |
| **Smokers** | | |
| - | 23 (77%) | 29 (64%) |
| No | 4 (13%) | 14 (31%) |
| Yes | 3 (10%) | 2 (4.4%) |
| **Disease state[3]** | | |
| Active | 22 (73%) | 23 (51%) |
| Remission | 8 (27%) | 22 (49%) |
| **CRP [mg/L]** | 0.23 (0.09, 0.94) | 0.07 (0.04, 0.13)*** |
| **5-aminosalicylic acid, Yes** | 29 (97%) | 44 (98%) |
| **Steroids, Yes** | 9 (30%) | 9 (20%) |
| **Azathioprine, Yes** | 17 (57%) | 13 (29%)* |
| **Biologics, Yes** | 12 (40%) | 11 (24%) |

* $p < 0.05$, ** $p < 0.01$, *** $p < 0.001$. Wilcoxon rank-sum test and Fisher's exact test were used for continuous and categorical variables.

[1] Data are presented as median (interquartile range) or n (%).

[2] A 150 µg/g cut-off was used to categorize participants into elevated or normal groups.

[3] The MAYO score cut-off of 1 and CD activity index (CDAI) score cut-off of 150 concerning CD and UC were used to reflect the disease activity in either active (i.e., mild, moderate, or severe conditions) or remission.

**Table S2. Clinical characteristics by disease activity state.**

| | **Active, N = 45**[1,2] | **Remission, N = 30**[1,2] |
|---|---|---|
| **Age at recruitment [years]** | 37 (26, 56) | 42 (26, 51) |
| **BMI [kg.m$^{-2}$]** | 22.4 (21.0, 24.7) | 22.0 (19.6, 23.7) |
| **Sex, females** | 13 (29%) | 14 (47%) |
| **Time since diagnosis [years]** | 1.05 (0.27, 1.90) | 2.18 (0.99, 5.15)* |
| **Smokers** | | |
| - | 33 (73%) | 19 (63%) |
| No | 10 (22%) | 8 (27%) |
| Yes | 2 (4%) | 3 (10%) |
| **Disease subtypes** | | |
| CD | 7 (16%) | 11 (37%) |
| UC | 38 (84%) | 19 (63%) |
| **Fecal calprotectin**[3] | | |
| Elevated | 22 (49%) | 8 (27%) |
| Normal | 23 (51%) | 22 (73%) |
| **CRP [mg/L]** | 0.09 (0.04, 0.31) | 0.10 (0.05, 0.27) |
| **5-aminosalicylic acid, Yes** | 44 (98%) | 29 (97%) |
| **Steroids, Yes** | 11 (24%) | 7 (23%) |
| **Azathioprine, Yes** | 17 (38%) | 13 (43%) |
| **Biologics, Yes** | 16 (36%) | 7 (23%) |

* $p < 0.05$. Wilcoxon rank-sum test and Fisher's exact test were used for continuous and categorical variables.

[1] Data are presented as median (interquartile range) or n (%).

[2] The MAYO score cut-off of 1 and CD activity index (CDAI) score cut-off of 150 concerning CD and UC were used to reflect the disease activity in either active (i.e., mild, moderate, or severe conditions) or remission.

[3] A 150 µg/g cut-off was used to categorize participants into elevated or normal groups.

**Table S3. Clinical characteristics by biologics status.**

| | Pre-, N = 10[1] | Post-, N = 13[1] |
|---|---|---|
| **Age at recruitment [years]** | 28 (25, 37) | 26 (24, 48) |
| **BMI [kg.m$^{-2}$]** | 23.33 (21.18, 25.30) | 22.70 (20.74, 25.10) |
| **Sex, females** | 2 (20%) | 5 (38%) |
| **Time since diagnosis [years]** | 0.92 (0.17, 4.04) | 1.74 (1.43, 2.12) |
| **Smokers** | | |
| - | 4 (40%) | 5 (38%) |
| No | 3 (30%) | 7 (54%) |
| Yes | 3 (30%) | 1 (7.7%) |
| **Disease subtypes** | | |
| CD | 3 (30%) | 5 (38%) |
| UC | 7 (70%) | 8 (62%) |
| **Disease state[2]** | | |
| Active | 8 (80%) | 8 (62%) |
| Remission | 2 (20%) | 5 (38%) |
| **Fecal calprotectin[3]** | | |
| Elevated | 7 (70%) | 5 (38%) |
| Normal | 3 (30%) | 8 (62%) |
| **CRP [mg/L]** | 0.19 (0.08, 0.31) | 0.04 (0.03, 0.09) |
| **5-aminosalicylic acid, Yes** | 10 (100%) | 12 (92%) |
| **Steroids, Yes** | 8 (80%) | 6 (46%) |
| **Azathioprine, Yes** | 7 (70%) | 7 (54%) |
| **Biologics, Yes** | 10 (100%) | 13 (100%) |

* $p < 0.05$. Wilcoxon rank-sum test and Fisher's exact test were used for continuous and categorical variables.

[1] Data are presented as median (interquartile range) or n (%).

[2] The MAYO score cut-off of 1 and CD activity index (CDAI) score cut-off of 150 concerning CD and UC were used to reflect the disease activity in either active (i.e., mild, moderate, or severe conditions) or remission.

[3] A 150 µg/g cut-off was used to categorize participants into elevated or normal groups.

98 **Table S4. Clinical characteristics by the need for treatment escalation.**

| Characteristic | No, N = 52[1] | Yes, N = 10[1] |
|---|---|---|
| **Age at recruitment [years]** | 43 (28, 58) | 28 (25, 37) |
| **BMI [kg.m$^{-2}$]** | 22.0 (20.7, 23.8) | 23.3 (21.2, 25.3) |
| **Sex, females** | 20 (38%) | 2 (20%) |
| **Time since diagnosis [years]** | 1.16 (0.37, 2.37) | 0.92 (0.17, 4.04) |
| **Smokers** | | |
| - | 43 (83%) | 4 (40%) |
| No | 8 (15%) | 3 (30%)* |
| Yes | 1 (1.9%) | 3 (30%)* |
| **Disease subtypes** | | |
| CD | 10 (19%) | 3 (30%) |
| UC | 42 (81%) | 7 (70%) |
| **Disease state[2]** | | |
| Active | 29 (56%) | 8 (80%) |
| Remission | 23 (44%) | 2 (20%) |
| **Fecal calprotectin[3]** | | |
| Elevated | 18 (35%) | 7 (70%) |
| Normal | 34 (65%) | 3 (30%) |
| **CRP [mg/L]** | 0.11 (0.05, 0.30) | 0.19 (0.08, 0.31) |
| **5-aminosalicylic acid, Yes** | 51 (98%) | 10 (100%) |
| **Steroids, Yes** | 4 (7.7%) | 8 (80%)*** |
| **Azathioprine, Yes** | 16 (31%) | 7 (70%)* |
| **Biologics, Yes** | 0 (0%) | 10 (100%)*** |

* $p < 0.05$, ** $p < 0.01$, *** $p < 0.001$. Wilcoxon rank-sum test and Fisher's exact test were used for continuous and categorical variables.

[1] Data are presented as median (interquartile range) or n (%).

[2] The MAYO score cut-off of 1 and CD activity index (CDAI) score cut-off of 150 concerning CD and UC were used to reflect the disease activity in either active (i.e., mild, moderate, or severe conditions) or remission.

[3] A 150 µg/g cut-off was used to categorize participants into elevated or normal groups.

99
